# Structural basis for targeting human cancer antigen STEAP1 with antibodies

**DOI:** 10.1101/2020.03.07.981829

**Authors:** Wout Oosterheert, Piet Gros

## Abstract

Six-transmembrane epithelial antigen of the prostate (STEAP1) is an integral membrane protein that is highly upregulated on the cell surface of several human cancers, making it a promising therapeutic target. It shares sequence homology with three enzymes (STEAP2-4) that catalyze the NADPH-dependent reduction of iron(III). However, STEAP1 lacks an intracellular NADPH-binding domain and does not exhibit cellular ferric-reductase activity. Thus, both the molecular function of STEAP1 and its role in cancer progression remain elusive. Here, we present a ~3.0 Å cryo-electron microscopy structure of trimeric human STEAP1 bound to three Fab-fragments of the clinically employed antibody mAb120.545. STEAP1 adopts a reductase-like conformation and interacts with the Fabs through its extracellular helices. Enzymatic assays in human cells reveal that STEAP1 promotes iron(III) reduction when fused to the intracellular NADPH-binding domain of its family member STEAP4, implicating STEAP1 as a functional ferric reductase in STEAP hetero-trimers. Our work provides a foundation for deciphering the molecular mechanisms of STEAP1 and will be instrumental in the design of new therapeutic strategies to target STEAP1 in cancer.

## Introduction

Since its discovery in 1999 as multi-span membrane protein highly expressed on prostate cancer cells (1), six-transmembrane epithelial antigen of the prostate (STEAP1) emerged as cancer antigen expressed in various human cancers, which include prostate, bladder, colorectal, lung, ovarian, breast carcinoma and Ewing sarcoma. Because its expression in physiological tissue is minimal and mainly confined to the prostate gland (2), STEAP1 represents a potentially attractive therapeutic tool as both cancer biomarker and as a target for anti-cancer therapies (2–4). Indeed, several strategies for targeting STEAP1 in cancer have been explored; in 2007, a study reported the production and characterization of two monoclonal antibodies (mAb120.545 and mAb92.30) that bind STEAP1 with nanomolar affinity on prostate cancer cells and inhibit the growth of prostate and bladder-tumour xenografts in mice (5). More recently, clinical studies employing humanized variants of mAb120.545 that target STEAP1 were conducted, including: *1)* a phase I trial using an antibody-drug-conjugate (termed DSTP3086S or Vandortuzumab Vedotin) to target prostate cancer (6–8), and *2)* a combined Phase I / Phase II trial for the PET-imaging of metastatic castration-resistant prostate cancer patients using Zr^89^ labelled antibody (termed [^89^Zr]Zr-DFO-MSTP2109A) (9–11). Besides antibody-based strategies, several *in vitro* and *in vivo* studies revealed that STEAP1-derived peptides are immunogenic and thus suitable for recognition by cytotoxic T lymphocytes (12–16), indicating that STEAP1 could represent a potential candidate for the development of anti-cancer vaccines (4, 17).

STEAP1 belongs to a protein family that comprises three metalloreductases (18, 19) (STEAP2-4, also known as STAMP1-3 (20–22)), which reduce iron(III) and copper(II) and are also associated with cancer progression (23–25). At the molecular level, the four STEAP proteins are predicted to adopt a common architecture with intracellular N- and C-termini, six transmembrane helices and a single heme-b prosthetic group bound in the transmembrane domain (TMD) (26). STEAP2-4 additionally contain an intracellular oxidoreductase domain (OxRD) that binds nicotinamide-adenine dinucleotide phosphate (NADPH) (27, 28). The ferric and cupric reductase mechanism of STEAP2-4 is defined by electron transfer from intracellular NADPH through membrane-embedded flavin-adenine dinucleotide (FAD) and heme cofactors to chelated metal-ion complexes at the membrane extracellular side (26, 29). Contrary to STEAP2-4, STEAP1 does not exhibit metalloreductase activity when over expressed on mammalian cells (19), suggesting that it may have a distinct, yet unidentified function. However, a recent study revealed that dithionite-reduced, purified STEAP1 retains heme and is capable of reducing metal-ion complexes and oxygen (30), indicating that the absence of a binding site for an electron-donating substrate like NADPH could explain the lack of reductase activity for STEAP1. It has been proposed that STEAP1 may have a functional role in hetero-oligomeric complexes with other STEAP paralogs (19, 30). In support, its expression often correlates with the expression of STEAP2 in cancers (17) and both proteins co-purify in detergent (30), suggesting they could form a functional complex. Further indications for a functional hetero-trimeric STEAP complex emerged from the recent cryo-EM structures of homo-trimeric human STEAP4 (29), which revealed a domain-swapped architecture, with the intracellular OxRD positioned beneath the TMD of the adjacent protomer. This arrangement supports a model in which the heme in STEAP1 receives electrons from NADPH bound to an adjacent STEAP2/3/4 subunit. However, the *in vivo* redox activity of STEAP1, in both the absence and presence of other STEAP paralogs, remains to be established. In addition, there are no high-resolution structures available to help distinguish a functional role for STEAP1 as metalloreductase or, as previously proposed, a potential channel or transporter protein (1, 2, 5, 31). Thus, despite that STEAP1 is a populous plasma-membrane component of many different types of cancer cells and hence a promising novel therapeutic target, its structure and function in both health and disease remain unknown.

Here, we present the cryo-EM structure of full-length, trimeric human STEAP1 bound to three Fab fragments of the therapeutically relevant mAb120.545. The Fabs dock on the extracellular helices of STEAP1 through an extensive polar interface. The TMD of STEAP1 resembles the architecture of the STEAP4-TMD, and surprisingly, exhibits cellular ferric reductase activity when fused to the NADPH-binding OxRD of STEAP4.

## Results

### Biochemical characterization of STEAP1

A previous pioneering study reported the biophysical and electrochemical characterization of N-terminally truncated rabbit STEAP1, purified from insect cells in lauryl maltose neopentyl glycol (LMNG) detergent (30). Our initial attempts to purify full-length, human STEAP1 from mammalian HEK-cells using a similar protocol were hampered by the loss of the non-covalently bound heme-b cofactor during the purification, suggesting that STEAP1 was not natively folded. We therefore screened several other detergents for the solubilization of STEAP1 and identified digitonin as a suitable replacement for LMNG. In digitonin, the purified protein retained its heme cofactor (Fig. 1A) and eluted as a monodisperse peak in size exclusion chromatography (SEC) experiments (Fig. 1B, D). In addition, thermostability assays revealed a melting temperature (T_M_) of ~55.5 °C for STEAP1 in digitonin (Fig. 1D, E), indicating that the protein was stable outside its native membrane environment. To assess if purified STEAP1 adopted a physiological-relevant conformation, we tested the binding of STEAP1 to the Fab fragment of monoclonal antibody mAb120.545, which exhibits 1 nM affinity for STEAP1 on cells (5). Size-exclusion chromatography assays revealed a smaller elution volume for STEAP1 when it was premixed with Fab120.545 (Fig. 1B), suggesting the formation of a complex, which was then confirmed by SDS page gel analysis of the eluted sample (Fig. 1C). Thus, the conformation of the STEAP1-epitope recognized by the Fab fragment on cells is preserved during the detergent solubilization and purification of STEAP1.

**Figure 1:**
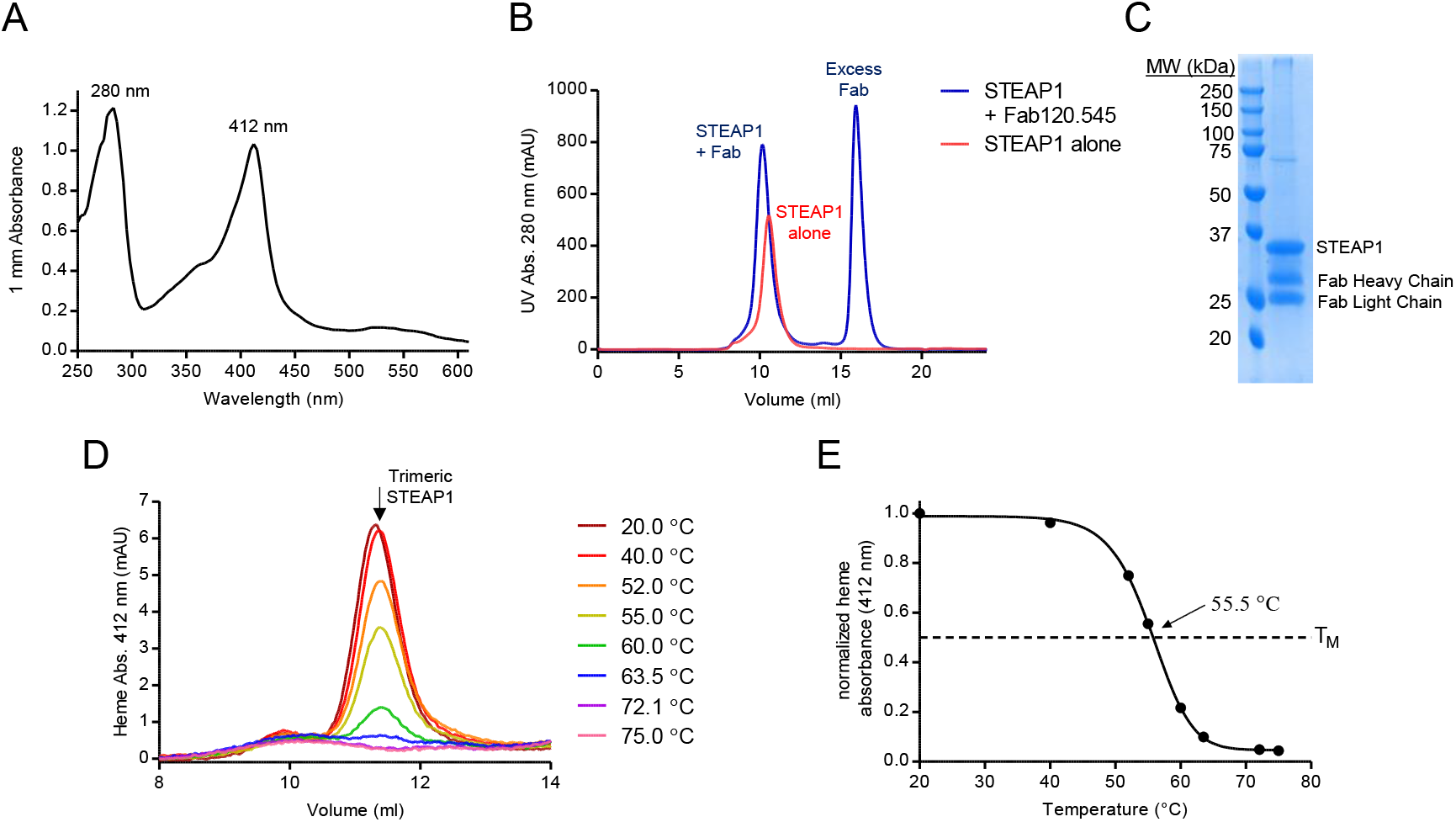
STEAP1 purification and stability. (A) UV-VIS spectrum of the purified STEAP1-Fab120.545 complex used for EM sample preparation. The protein and heme-absorbance peaks exhibit a maximum at 280 and 412 nm, respectively. (B) Size-exclusion chromatography elution profile of STEAP1 in the absence (red) or presence (blue) of an excess Fab120.545. (C) SDS page gel of the purified STEAP1-Fab120.545 complex. (D) Heme absorbance size-exclusion chromatography elution profiles of STEAP1 after 10 min incubation at several temperatures. (E) Melting curve for digitonin-purified STEAP1, generated by using the peak maxima from panel D. STEAP1 exhibits a T_m_ of 55.5 °C.

### Cryo-EM structure determination

To gain insights into the molecular architecture of STEAP1, we set out to obtain a structural model of the protein using single-particle cryo-EM. Full-length, trimeric STEAP1 proved to be a challenging sample for EM due to its small size (<120 kDa) and the absence of folded domains protruding from the membrane region. To create a larger particle with more extramembrane features to facilitate EM-image processing, we opted to determine the structure of STEAP1 purified in complex with Fab120.545. The complementarity-determining regions (CDRs) of Fab120.545 are identical to those present in STEAP1-antibodies used in clinical trials (Fig. S1), indicating that the structure of the STEAP1-Fab120.545 complex could also be instrumental in engineering antibodies and other molecules that target STEAP1 in cancer. Micrographs collected on a 200 kV Talos Arctica microscope showed non-aggregated particles distributed in vitreous ice (Fig. S2A). Subsequent 2D classification experiments yielded class averages with clear secondary structure elements and furthermore revealed that more than one Fab fragment is bound to micelle-embedded STEAP1 (Fig. 2A, Fig. S2B). Image processing in Relion (32) finally resulted in a reconstructed cryo-EM-density map at ~3.0 Å resolution (Fig. 2B, Fig. S2C-F). The map displayed well-defined sidechain density for the TMD of STEAP1 and the variable regions of the Fab (Fig. S3). The model for STEAP1 was built with the TMD of STEAP4 as the template (29) (pdb 6HCY), whereas the starting model for the Fab was generated through the PIGS homology server (33). The refined structure has acceptable stereochemistry and exhibits high correlation to the cryo-EM density map within the determined resolution (Fig. S3G, H, Table 1).

**Figure 2:**
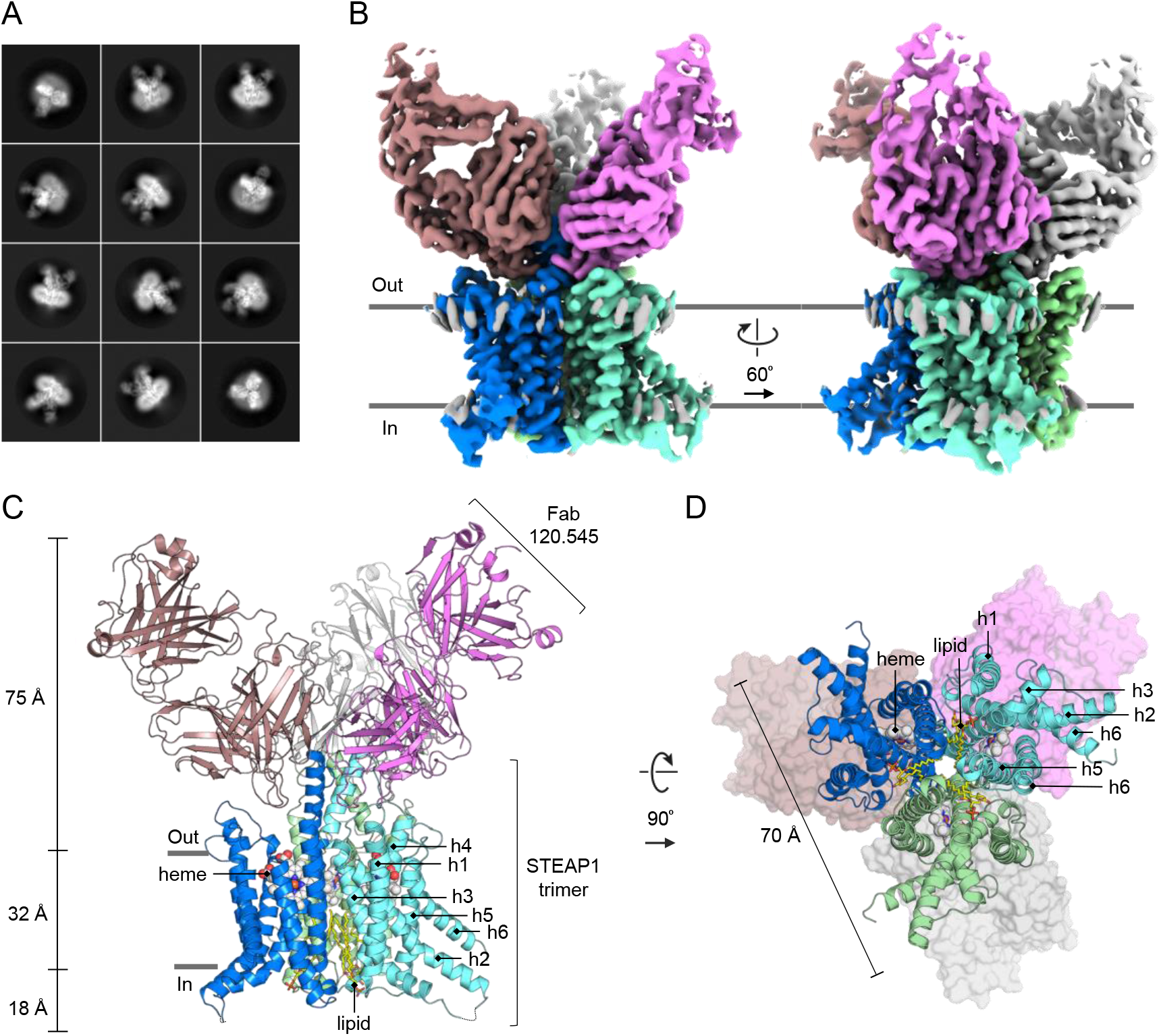
Cryo-EM structure of the STEAP1-Fab120.545 complex. (A) Exemplary 2D class averages. The box size corresponds to 300×300 pixels (309×309 Å). More 2D class averages are shown in Fig. S3. (B) Two orientations of the unsharpened cryo-EM density map at 3.0-Å resolution, colored by chain (STEAP1) or molecule (Fabs). (C, D) Atomic model of the STEAP1-Fab120.545 complex as viewed parallel to the membrane as a sideview C) or orthogonal to the membrane from the cytoplasmic side (D). In panel D, the Fabs are shown as surface.

**Table 1:**
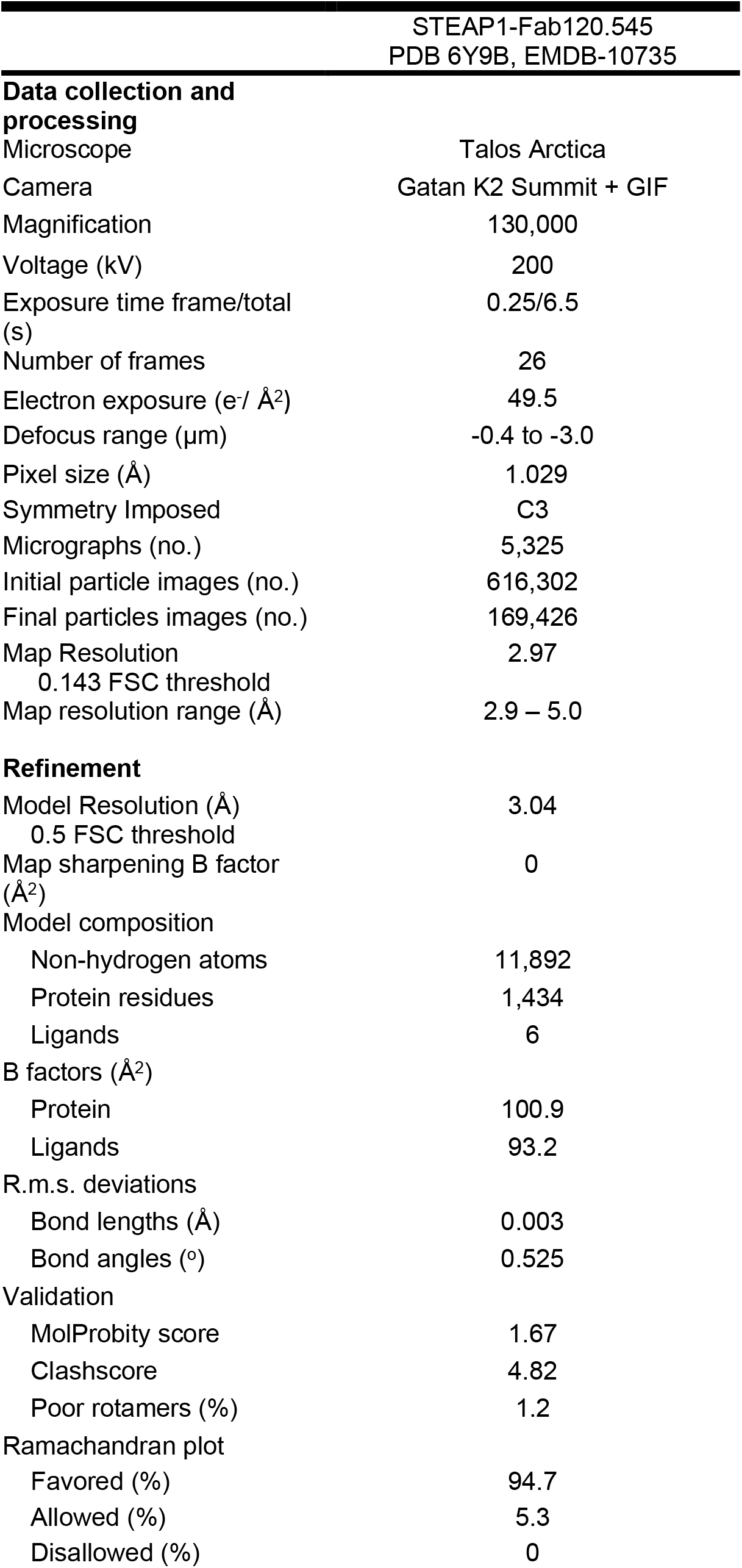
Cryo-EM data collection, refinement and validation statistics.

### Architecture

The cryo-EM structure reveals a one-to-one stoichiometry of the STEAP1-Fab120.545 complex, with three STEAP1 protomers interacting with three Fab molecules (Fig. 2B-D). The Fabs bind at the extracellular region of STEAP1, consistent with the observation that the antibody targets STEAP1 expressed on intact cancer cells. The intracellular loops of STEAP1 extend ~18 Å from the membrane region into the cytoplasm, whereas the Fabs protrude up to ~75 Å into the extracellular space. STEAP1 adopts a trimeric arrangement that is strikingly similar to the one of its family member STEAP4 (41% amino-acid sequence identity, rmsd 0.8 Å for 640 Cα atoms, Fig. 3). Each STEAP1 subunit comprises six membrane-spanning α-helices (h1 – h6) that define the TMD of the protein. A single b-type heme cofactor is sandwiched between helices h2 – h5 at the extracellular membrane leaflet (Fig. S3B). Strictly conserved histidine residues H175 and H268 coordinate the central iron moiety of the heme prosthetic group, thereby resembling the hexa-coordinated heme arrangement of STEAP4. At the intracellular membrane leaflet side of the TMD, we observed weak density not corresponding to any protein residues. An overlay with the structure of STEAP4 revealed that the observed density overlaps with the flavin ring of the FAD-binding site in STEAP4 (Fig. S3D). The FAD-interacting residues in the TMD of STEAP3 and 4 are conserved in STEAP1 and the STEAP1-Fab120.545 cryo-EM sample was supplemented with 1 mM FAD before grid freezing. However, STEAP3 and 4 additionally interact with the adenine-moiety of FAD via their intracellular OxRD (26, 29) (Fig. 3B, Fig. S3D), which is missing in STEAP1. In line with this, STEAP3 and 4 exhibit a low micromolar affinity for FAD (Kd = ~1 μM) (26, 29), whereas the affinity of STEAP1 for FAD is much weaker (Kd = 34 μM) (30). The cryo-EM density in this region could therefore correspond to a loosely bound FAD cofactor, although the weak density does not allow for modelling the complete cofactor. Instead of an OxRD of ~175 amino acids in length, STEAP1 contains a 69 residue N-terminal intracellular tail with no predicted domain architecture. Indeed, we observed no density for the first 65 intracellular-amino acids of STEAP1, indicating that these residues are flexible, which is consistent with in-silico disorder predictions using RONN (34). Thus, contrary to other STEAP family members, the homo-trimeric human STEAP1 structure does not harbor a folded N-terminal intracellular domain.

**Figure 3:**
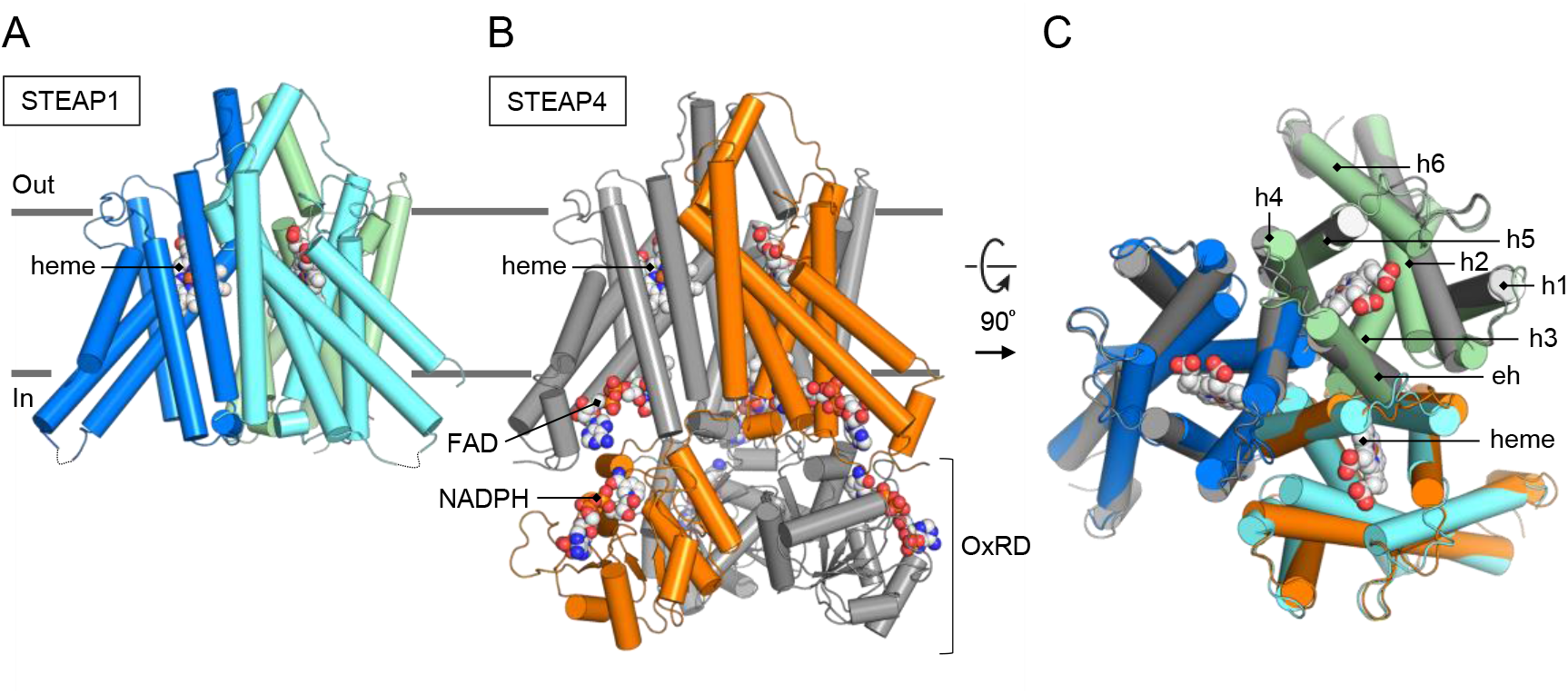
Structural similarities between human STEAP1 and STEAP4. (A, B) Aligned models (rmsd 0.8 Å for 640 Cα atoms) of STEAP1 (A) and STEAP4 (B) shown parallel to the membrane. (C) Overlay of STEAP1 and the TMD of STEAP4 (residues 195 – 454) shown orthogonal to the membrane from the extracellular side. Overlapping membrane helices are annotated.

### STEAP1-Fab120.545 interface

We next analyzed the interface between STEAP1 and Fab120.545. The epitope recognized by the Fab spans the first and second extracellular regions (EC1 and EC2, respectively) of STEAP1, which bridge membrane helices h1-h2 and h3-h4. The total interface formed between the STEAP1 trimer and three Fabs comprises ~5,730 Å^2^ of buried surface area and is arranged so that a single Fab molecule interacts with all three STEAP1 protomers. The interface is stabilized by a network of polar and hydrophobic interactions and can largely be described by two ‘interaction hotspots’ on the EC2 of STEAP1 (Fig. 4A, B). The first hotspot involves the extracellular helix of STEAP1 (residues 186 – 201), which extends from membrane helix h3. Y190, N194, W195 and Q198 interact with Fab heavy and light chain residues Y102_H_, Y104_H_, Y108_H_, Y31_L_ and S33_L_ (Fab chain identifier in subscript) (Fig. 4A). The carboxyl group of S33_L_ furthermore forms a hydrogen bond with the sidechain of Y107 of the EC1, whereas Y104_H_ bridges two STEAP1 protomers by interacting with both Y190 and W195 from different chains. The second hotspot comprises residues Q201, Q202, N203, Q205 and D206, which reside in the loop that connects the extracellular helix of STEAP1 to membrane helix h4. Hotspot-two consists of numerous polar interactions, including a salt bridge between D206 and R32_L_. Other Fab residues involved in binding to hotspot-two are Y51_H_, S57_H_, T58_H_, S59_H_, Q27_L_, S33_L_,N99_L_ and Y100_L_ (Fig. 4B). Besides the Fab residues in close vicinity (<4 Å) to STEAP1, we identified three aspartate residues (D103_H_, D105_H_, D106_H_) in the Fab heavy chain that orient towards the basic amino-acid ring above the heme (Fig. 4C). These aspartates are at least 4.5 Å separated from any STEAP1-residues and could participate in long-range electrostatic interactions with the basic ring of STEAP1. Interestingly, in STEAP4 the corresponding basic amino-acids comprise the substrate-binding site (Fig. 4D). To experimentally verify the STEAP1-Fab interface observed in the structure, we generated several mutants of Fab120.545 and tested their ability to bind purified STEAP1 using size-exclusion chromatography assays (Fig. S4). Mutants R32_L_E and N99_L_D were designed to create charge repulsions in hotspot-2 between STEAP1 and Fab120.545. As expected, we did not observe binding events for these two mutants (Fig. S4A-C). Intriguingly, Fab mutants D103_H_N,D105_H_N,D106_H_N (Fab-NNN) and D103_H_A,D105_H_A,D106_H_A (Fab-AAA) similarly did not interact with purified STEAP1 (Fig. S4A, D, E), indicating that the long-range electrostatic interactions between the three Fab-aspartates and STEAP1 are essential for maintaining a high-affinity antibody-antigen complex.

**Figure 4:**
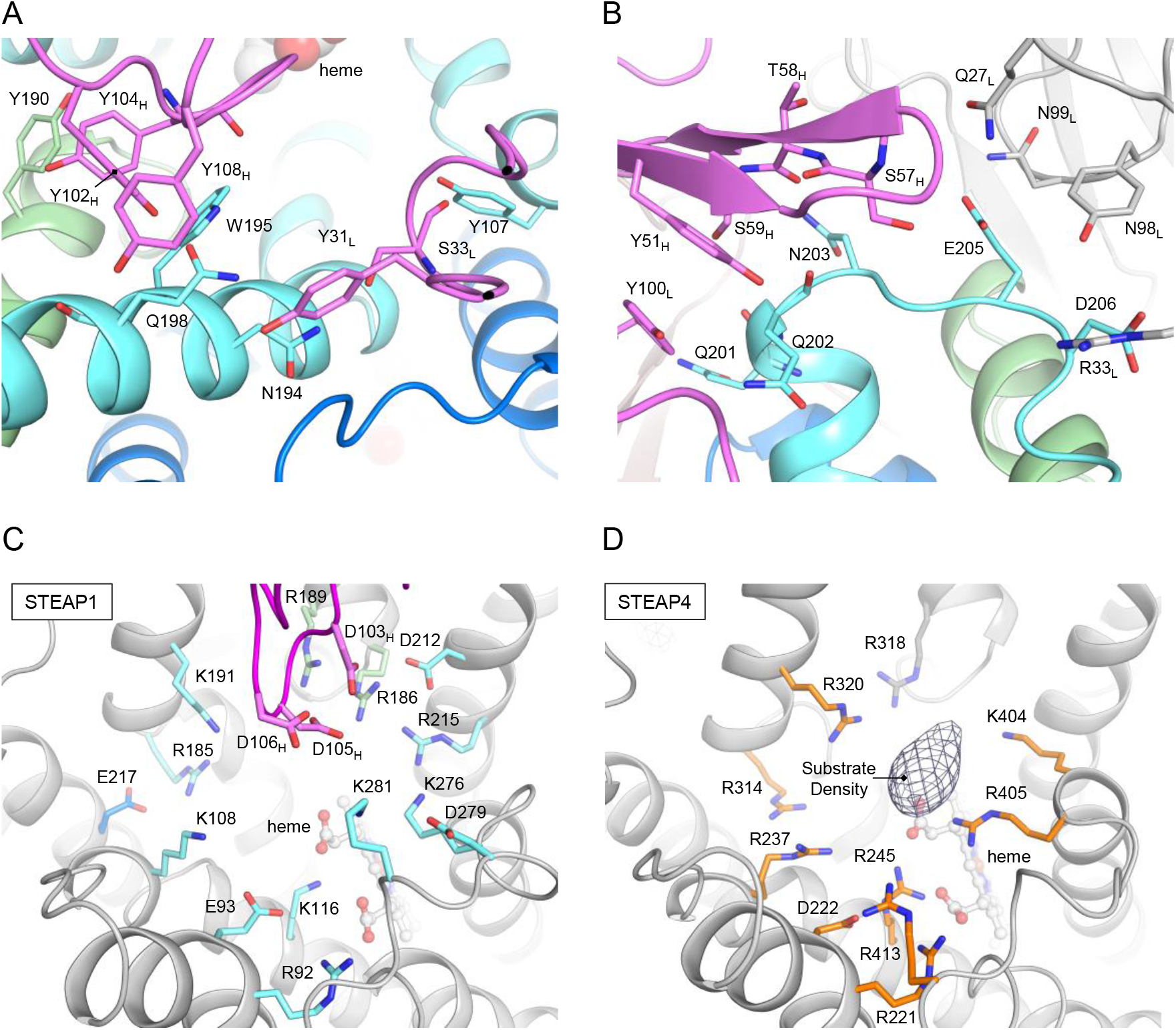
Fab120.545-binding at the putative substrate binding site. (A, B) Interactions between STEAP1 and Fab120.545 at (A) hotspot 1, corresponding to the extracellular helix of STEAP1 and (B) hotspot 2, the loop between the extracellular helix and membrane helix h4. Amino acids backbones are only shown in sticks when they contribute to the interface. (C) Amino acid environment above the heme in the STEAP1-Fab structure. All charged residues are shown as sticks. The Fab light chain is omitted from the figure for clarity. STEAP1 helices are colored grey for clarity. (D) Amino acid-environment above the heme in the STEAP4 structure (pdb 6HCY, EMDB-0199). All charged residues are shown as sticks. The difference density (taken from EMDB-0199) that corresponds to substrate Fe(III)-NTA is depicted in mesh.

### Generation of a functional STEAP4/1 fusion protein

The reductase-like architecture of STEAP1 (Fig. 3) and its heme-redox potential of −114 to −118 mV (30), indicate that the protein could be functional in reducing metal-ion complexes *in vivo*. However, to the best of our knowledge there is currently no experimental data available that shows that STEAP1 is capable of reducing ferric iron in a physiological setting. Although STEAP1 may form relevant heterotrimeric complexes with other STEAP homologs (30), co-expressions of different STEAP family members will likely result in a mixed population of homo- and heterotrimers, making cellular ferric-reductase experiments difficult to interpret. To overcome these hurdles and to assess if the TMD of STEAP1 can direct electron transport across mammalian cell membranes, we aimed to design a construct in which the STEAP1-TMD is fused to the intracellular OxRD of another STEAP homolog. To this end, a sequence alignment of all human STEAP proteins revealed that STEAP4 and STEAP1 share a common three-amino acid LFP-motif at the start of membrane helix h1. Additionally, an overlay of their cryo-EM structures did not show any obvious clashes between the OxRD of STEAP4 and the TMD of STEAP1. Thus, we generated a construct that spans residues M1 – Q195 of STEAP4, the shared LFP motif and residues Q69 – L339 of STEAP1, which we termed STEAP4/1_chimera_ (Fig. 5A, B). We then expressed STEAP4/1_chimera_ in HEK293 cells and compared its cellular reductase activity to cells expressing STEAP1 or STEAP4, using the physiologically relevant ferric-citrate as a substrate. Consistent with a previous study (19), over-expression of STEAP1 did not result in measurable cell-surface ferric-reductase activity compared to the empty-vector control, whereas cells expressing STEAP4 reduced ~57 pmol Fe^3+^ per minute per well (Fig. 5C). Intriguingly, the STEAP4/1_chimera_ expressing cells also showed a highly significant reductase activity of ~43 pmol Fe^3+^ per minute per well (Fig. 5C). To verify that the observed activity was dependent on transmembrane-electron transport through the TMD of STEAP1, we additionally tested STEAP4/1_chimera_ mutants R161E and H175A (STEAP1 numbering) in which, respectively, the FAD and heme-binding sites in the TMD are abolished. Cells expressing STEAP4/1_chimera_-R161E and STEAP4/1_chimera_-H175A did not exhibit any ferric reductase activity (Fig. 5C), indicating that the STEAP1-TMD of the chimera is indeed crucial for cell-surface iron reduction. Confocal microscopy experiments subsequently confirmed that all expressed proteins except for STEAP4/1_chimera_-H175A localize to the plasma membrane (Fig. S5). Taken together, our cell-based experiments reveal that STEAP1 adopts a conformation that facilitates transmembrane-electron transport to reduce ferric citrate at the membrane extracellular side. Therefore, the lack of reductase activity of the protein can be explained by the absence of a binding site for an electron-donating substrate in homo-trimeric STEAP1.

**Figure 5:**
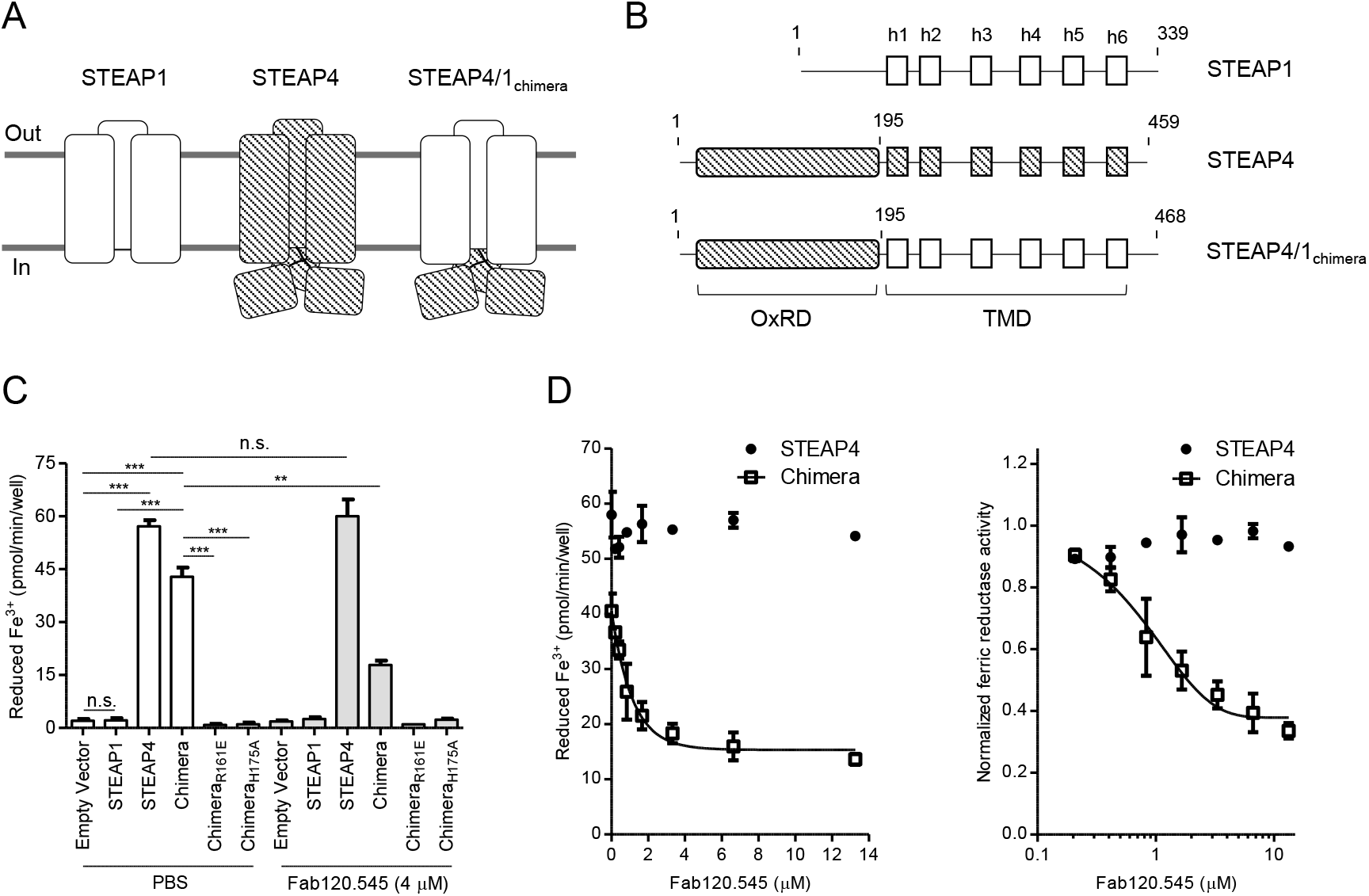
Cellular ferric-reductase activity of STEAP protein variants. (A) Schematic topological representation of natural occurring STEAP1 and STEAP4 and the designed fusion protein STEAP4/1_chimera_. Domains of STEAP1 are shown in solid white, whereas STEAP4 domains are depicted in diagonal stripes. (B) Domain composition of STEAP1, STEAP4 and STEAP4/1_chimera_ constructs used for cellular ferric-reductase experiments. (C) Cell-surface ferric-reductase activity of STEAP protein variants supplemented with PBS or with 4 μM Fab120.545. The experiment was performed in triplicate, error bars represent the standard deviation. (D) Cell-surface ferric-reductase activity of STEAP4 and STEAP4/1_chimera_ while varying the concentration of added Fab120.545. The STEAP4/1_chimera_ datapoints are fitted with a one-phase decay curve. Error bars represent the standard deviation between triplicate experiment. The right panel shows the same data normalized to an activity of 1 at 0 μM Fab120.545, with a logarithmic x-axis.

### Fab120.545 inhibits ferric reductase activity of STEAP4/1_chimera_

Because Fab120.545 binds in close proximity to the putative substrate-binding site in STEAP1 (Fig. 4C, D), we assessed if Fab binding could influence the ferric reductase activity of STEAP4/1_chimera_. Indeed, the addition of 4 μM Fab120.545 to cells expressing STEAP4/1_chimera_ led to a significant decrease in iron-citrate reduction from ~43 to ~18 pmol Fe^3+^ per minute per well (Fig. 5C). Conversely, the activity of STEAP4 was unchanged, indicating that the Fab specifically recognizes an extracellular epitope of STEAP1. We further characterized the inhibitory properties of the Fab by incubating cells expressing STEAP4 or STEAP4/1_chimera_ with different Fab concentrations. This revealed a Fab-concentration dependent effect on the inhibition of the ferric reductase activity of STEAP4/1_chimera_, while the amount of iron(III) reduced by STEAP4 remained unaltered over the entire concentration range tested (Fig. 5D).

## Discussion

Based on its amino-acid sequence and subcellular localization, STEAP1 was previously predicted to function as a channel or transporter protein (1, 2, 5, 31). The cryo-EM structure of STEAP1 bound to antibody-fragment Fab120.545 revealed a trimeric arrangement similar to the one of its family member STEAP4 (Fig. 3), showing no obvious structural features of ion channels or transporters. Instead, the *1)* strictly conserved FAD-binding residues and putative loosely-bound FAD at the inner-membrane leaflet of the TMD (Fig. S3D), *2)* heme-binding site at the outer-membrane leaflet (Fig. S3B) and *3)* basic amino-acid ring above the heme (Fig. 4C), all imply that STEAP1 may function as a transmembrane oxidoreductase. In our previous work, we showed that substrate iron(III), complexed to a negatively-charged chelator like citrate, binds in the basic ring of STEAP4 (Fig. 4D) and we proposed that this ring of positive amino acids may polarize the iron(III)-chelator complex to facilitate the iron-reduction reaction (29). The presence of a comparable positively charged ring (Fig. 4C, D) indicates that STEAP1 harbors a similar protein environment for the reduction of iron(III).

We investigated whether the TMD of STEAP1 is capable of directing transmembrane-electron transport by generating a fusion construct between the intracellular region of STEAP4 and the TMD of STEAP1, termed STEAP4/1_chimera_ (Fig. 5A, B). Cell-surface expressed STEAP4/1_chimera_ catalyzed the reduction of iron-citrate, providing evidence that STEAP1 is a functional reductase, albeit only when a binding site for intracellular electron-donating substrate NADPH is available (Fig. 5C, D). The lack of enzymatic activity of STEAP4/1_chimera_-R161E (Fig. 5C), that localizes to the plasma membrane in HEK-cells (Fig. S5D), confirmed that the TMD of STEAP1 enables transmembrane-electron transport. We also showed that STEAP4/1_chimera_-H175A exhibited no significant ferric-reductase activity (Fig. 5C). However, fluorescence microscopy experiments revealed that this mutant resides almost exclusively in intracellular compartments (Fig. S5E), suggesting that the protein misfolds when the heme-cofactor binding site is abolished.

The addition of Fab120.545 resulted in a concentration-dependent decrease of ferric reductase activity of STEAP4/1_chimera_-expressing cells, but not of STEAP4-expressing-cells (Fig. 5C, D). This demonstrates that the Fab likely does not interact with STEAP4, which can be explained by the observation that the glycan on STEAP4-residue N323 would clash with the Fab light chain. In contrast, the residue at the equivalent position in STEAP1 (N194) is not glycosylated and forms a hydrogen bond with S33_L_ (Fig. 4A).

How does Fab120.545 inhibit the cell-surface ferric reductase activity of STEAP4/1_chimera_? The conformations of the extracellular regions of STEAP4 and Fab-bound STEAP1 are similar (Fig. 3), so the binding of Fab120.545 is not expected to induce large conformational changes in STEAP1. Instead, the Fab partially blocks access to the putative substrate-binding site in the basic ring above the heme (Fig. 4C). Alternatively, Fab residues D103_H_, D105_H_ and D106_H_ might neutralize the positive-charged substrate-binding site and thereby prevent substrate polarization. However, mutagenesis of these aspartates to either asparagines (Fab-NNN) or alanines (Fab-AAA) resulted in a loss of binding of Fab120.545 to purified STEAP1 (Fig. S4A, D, E), hence this hypothesis could not be tested. Nevertheless, our results suggest that antibodies can be employed as tools to inhibit the ferric-reductase activity of STEAP enzymes. Additionally, the 3:3 STEAP1:Fab120.545 stoichiometry (Fig. 2B, C) indicates that full-length antibodies may crosslink STEAP1 trimers into higher-order assemblies on cell membranes. A similar antibody-induced crosslinking mechanism has recently been reported for the therapeutic antibody rituximab binding to its dimeric target protein CD20 (35).

In conclusion, the study presented here describes the first structure-function analysis of the human cancer antigen STEAP1. Our results support a model in which STEAP1 forms heteromeric assemblies with partner proteins that recruit and orient intracellular electron-donating substrates towards the TMD of STEAP1, enabling transmembrane-electron transport and the reduction of extracellular metal-ion complexes. This model warrants further investigations into the physiological function of STEAP1; for example, the incorporation of STEAP1 into STEAP heterotrimers might moderate iron(III)-reduction rates locally and thereby prevent deleterious reactions associated with iron overload. We therefore envision that it will be of great interest to focus future research endeavors on these putative assemblies of STEAP1 with STEAP2-4 family members and other unidentified accessory proteins in relevant cancer tissues. Ultimately, understanding the molecular principles that underly the function of STEAP1 will guide the design of anti-STEAP1 focused cancer therapies, thereby exploiting the protein’s high expression in cancer and minimal presence in healthy cells.

## Methods

### Chemicals

All chemicals were purchased from Sigma-Aldrich unless specified otherwise.

### Constructs

Codon-optimized DNA for mammalian cell expression encoding for human STEAP1 and STEAP4 was purchased from Geneart. The full-length STEAP1 construct used for structure determination was cloned in a pUPE expression vector (U-Protein Express BV) with C-terminal Strep3 tag. The STEAP4/1_chimera_ construct was generated through Gibson-assembly cloning (NEB). For functional assays in HEK-cells, all constructs were cloned in a pUPE expression vector with a C-terminal GFP-Strep3 tag with a TEV-protease site. Mutagenesis of STEAP construct was performed using the Q5 Site-Directed Mutagenesis Kit (NEB). The primers used in this study are listed in Table S1. In the manuscript, the amino-acid numbering of STEAP1 was used for mutations introduced in the STEAP4/1_chimera_ (R161E and H175A), because the mutated amino acids reside in the STEAP1-domain of the chimera. These residues correspond to R290 and H304 in both STEAP4/1_chimera_ and STEAP4.

### Protein expression and purification

The protein production protocol was adapted from the previously described protocol for STEAP4 (29). Full-length STEAP1 with a C-terminal Strep3 tag was expressed in HEK293 GNTI^−^ suspension cells (provided by U-Protein Express BV). Cells were grown at 37 °C for ~96h. All subsequent steps were performed at 4 °C, unless stated otherwise. After harvesting, cells were washed in PBS buffer and solubilized in lysis buffer containing 50 mM Tris pH 7.8, 250 mM NaCl, 0.7% (w/v) digitonin (Calbiochem), 0.3% (w/v) n-Dodecyl-β-D-Maltoside (DDM, Anatrace), 0.06% (w/v) Cholesteryl hemi-succinate (CHS) and protease inhibitor cocktail (Roche) for 2 – 3 hours. The sample was then subjected to ultracentrifugation at 100,000g for 45 min to remove insoluble membranes and cell debris. The supernatant was incubated with Streptactin resin (GE Healthcare) for 2h and the resin was washed with 20 column values of buffer A (50 mM Tris pH 7.8, 250 mM NaCl, 0.08% digitonin). Protein was subsequently eluted with buffer A supplemented with 3.5 mM desthiobiotin. STEAP1-containing fractions (which exhibited a red color due to the presence of the heme cofactor) were concentrated by a 100 kDa MW-cutoff concentrator device (Amicon) to ~1.6 mg/ml. Subsequently, 330 μl of STEAP1 was mixed with a large excess of Fab-fragment 120.545 (145 μl at 9.7 mg/ml in PBS) . After an hour of incubation, the STEAP1-Fab mixture was injected on a Superdex200 increase column (GE Healthcare) pre-equilibrated in 20 mM Tris pH 7.8, 200 mM NaCl, 0.08% (w/v) digitonin. Fractions containing the STEAP1-Fab complex were concentrated to a final concentration of ~5.0 mg/ml. Sample purity was assessed with SDS PAGE gel analysis and analytical size-exclusion chromatography.

### Grid preparation

Concentrated STEAP1-Fab120.545 (5.0 mg/ml) was incubated with 1 mM FAD on ice for one hour before grid freezing. 2.8 μl of sample was pipetted onto glow-discharged R1.2/1.3 200 mesh Au holey carbon grids (Quantifoil) and then plunge-frozen in a liquid ethane/propane mixture with a Vitrobot Mark IV (Thermo Fisher Scientific), blotting with force 0 for 4 seconds at 20 °C.

### EM data collection

Movie collection was performed on a 200 kV Talos Arctica microscope (Thermo Fisher Scientific) equipped with a K2 summit detector (Gatan) and a post column 20 eV energy filter. Using EPU (Thermo Fisher Scientific) in superresolution/counting mode (pixel size 0.514 Å), movies were collected for 6.5 s in 26 frames with a dose of 1.905 e^−^/Å^2^/frame (measured in an empty hole without ice), corresponding to a total-electron exposure of 49.5 e^−^/Å^2^. Defocus values for collection in EPU were set between −1 and −3 μm, but varied between −0.4 and −3.5 μm during data collection.

### Image processing

5,352 movies were imported in the Relion3.0 pipeline. The movies recorded in super-resolution mode were binned 2× (resulting pixel size 1.03 Å) and motion corrected using UCSF MotionCor2 (36), followed by CTF estimation using GCTF (37). 695 movies were subsequently discarded based on their poor CTF spectra, resulting in 4,657 movies (87% of total) for further processing. 1,791 particles were picked manually and 2D classified. The generated classes were used as templates for autopicking in Relion (38), resulting in 616,302 particles. The picked particles were 3D classified into 6 classes with no symmetry applied. The particles belonging to the class with most protein-like features (263,939 particles) were then subjected to CTF refinement and Bayesian polishing, followed by 3D classification without image alignment into 3 classes. The highest populated class (172,724 particles) showed clear amino-acid side chain features. Particles were then CTF-refined for a second time and 431 junk particles were removed through 2D classifications. 3D Auto refinement (with C3 symmetry applied) of the remaining 172,293 particles yielded a map at a global resolution of 3.8 Å based on the gold-standard FSC = 0.143 criterion. A postprocessing step in which the constant region of the Fab was masked out, improved the resolution to 3.5 Å. Following the release of Relion version 3.1beta with high-order aberration and anisotropic magnification estimation (39), we performed four additional rounds of CTF refinement and Bayesian polishing. This iterative process was followed by a 3D classification without image alignment into 3 classes, removing 2,867 particles. The final 169,426 particles were 3D Auto Refined (C3 symmetry) and subjected to a postprocessing step, improving the map resolution to 3.0 Å (3.3 Å without masking), corresponding to ~1.44× Nyquist frequency.

### Model building and refinement

To build the model for STEAP1, the TMD structure of human STEAP4 (residues 196 – 454) was rigid-body fitted into the cryo-EM map. For one chain, all STEAP4 residues were changed to the corresponding STEAP1 residue using the ‘mutate residue range’ option in Coot (40), after which the model was manually inspected, adjusted and refined in Coot. The model of this chain was copied and fitted in the density of the other two subunits. The starting model for the variable regions of Fab120.545 was obtained through the PIGS homology server (33) by uploading the heavy and light chain sequences. This model was rigid-body fitted in the cryo-EM map and the CDR regions were manually built in Coot. Then, the STEAP1-Fab120.545 model was iteratively refined using Coot (manually) and Phenix real-space refine (41) with geometric restraints and non-crystallographic symmetry (NCS) constraints. Final refinements were performed using the non-sharpened cryo-EM map, in which the constant region of the Fab was masked out. The non-sharpened map revealed sufficient side-chain detail for modelling. The final model uploaded to the pdb comprises residues 67 – 312 of STEAP1, residues 1 – 112 of the Fab120.545 light chain and residues 2 – 122 of the Fab120.545 heavy chain.

### Thermostability assays

Thermostability assays were performed as previously reported (29, 42, 43). Aliquots of purified STEAP1 (in 20 mM Tris pH 7.8, 200 mM NaCl, 0.08% digitonin) were heated over a range of temperatures (20 – 75 °C) in a thermocycler for 10 min, cooled down and centrifuged to remove aggregates. The supernatant was subsequently injected on a Superdex200 10/300 increase column equilibrated in 20 mM Tris pH 7.8, 200 mM NaCl, 0.08% digitonin, connected to a HPLC system (Shimadzu). The heme absorbance of STEAP1 was monitored at 412 nm using a SPD-20A UV-Vis detector. In order to determine the melting temperature (T_m_), peak maxima were normalized to the sample incubated at 20 °C and fitted to a dose response equation using GraphPad Prism 5.

### SEC-binding assays with Fab variants

All Fab variants were diluted to a concentration 0.5 mg/ml in PBS supplemented with 0.08% digitonin. 25 μl of purified STEAP1 (0.28 mg/ml in 20 mM Tris pH 7.8, 200 mM NaCl, 0.08% digitonin) was mixed with 35 μl PBS + digitonin or a ~2 fold molar excess of Fab (35 μl at 0.5 mg/ml). After incubation for several hours, the mixtures were injected on a Superdex200 3.2/300 increase column equilibrated in 20 mM Tris pH 7.8, 200 mM NaCl, 0.08% digitonin, connected to a HPLC system (Shimadzu). The heme absorbance of STEAP1 was monitored at 412 nm using a SPD-20A UV-Vis detector, whereas the Tryptophan fluorescence (ex 275, em 354) emitted by both STEAP1 and Fab variants was detected using RF-20A xs detector. STEAP1-Fab complex formation was assessed by comparing the peak elution profiles of mixtures with the profiles of individually injected proteins.

### Ferric-reductase assays

HEK293 GNTI^−^ suspension cells (U-Protein Express BV) were transfected with GFP-tagged STEAP constructs. ~96 hours after transfection, cells were washed in PBS, resuspended in iron uptake buffer (25 mM MES, 25 mM MOPS pH 7.0, 140 mM NaCl, 5.4 mM KCl, 1.8 mM CaCl_2_, 0.8 mM MgCl_2_, 5 mM glucose, 400 μM ferrozine) and pipetted in a 96-well plate (~5*10^4^ cells per well). Experiments were started by the addition of ferric citrate (Fisher Scientific) to each well (200 μM final concentration). The assay was performed in the dark at 37 °C for 35 minutes. Fe^2+^-ferrozine formation was monitored using a Model 680 microplate reader (Biorad) at 550 nm. The formed Fe^2+^ was quantified using a standard curve that was generated as described (44). To assess the effect of Fab120.545 on the ferric reductase activity of the STEAP variants, cells were incubated with PBS or a concentration range of Fab120.545 for 20 minutes prior to the addition of ferric citrate. Experiments were performed in triplicate by diluting cell stocks three separate times. Error bars represent the standard-deviation. Experiments in which activities were compared, were carried out in parallel in the same 96-well plate. All statistical analyses were performed in Graphpad Prism 5.0. The ferric reductase-activities of cells expressing different STEAP variants were compared for statistical significance using unpaired t-tests, whereas paired t-tests were employed to compare the same population of cells with or without the addition of Fab120.545.

### Fluorescence Microscopy

HEK293 GNTI^−^ suspension cells (U-Protein Express BV) were transfected with GFP-tagged STEAP constructs. ~96 hours after transfection, cells were washed in PBS and then imaged for GFP (ex 488 nm, em 509 nm) using a Corrsight spinning-disk confocal microscope (FEI) at a magnification of 40x at 20 °C.

### Data availability

Data supporting the findings of this manuscript are available from the corresponding authors upon reasonable request. The relevant cryo-EM density maps of the STEAP1-Fab120.545 complex have been deposited under accession number EMDB-10735. This deposition includes unfiltered-half maps, non-sharpened unmasked maps and sharpened masked maps. Model coordinates of the structure have been deposited in the Protein Data Bank under accession number 6Y9B.

## Acknowledgements

The Fab120.545 fragments were a kind gift from Genmab BV and were produced upon our request. We furthermore gratefully thank W. Hemrika (U-Protein Express BV) for HEK cell cultures; L.S. van Bezouwen, S.C. Howes and the Utrecht EM-square (C.T.W.M. Schneijdenberg and J.D. Meeldijk) for pivotal electron-microscope assistance; A. M. Liaci for help with Gibson-assembly cloning; J. Granneman for initial construct cloning; D. El Mazouni for help with confocal microscopy; T.H. Brondijk and R.M. v.d. Bos for proofreading of the manuscript. This work has been supported by the Netherlands Organization for Scientific Research (NWO), Fund NCI Technology Area (project no. 731.015.201).

## Author contributions

W.O and P.G designed the project. W.O carried out all experiments, analyzed the data and wrote the manuscript under the guidance of P.G.

