## Supplemental Figures S1 - S5, Table S1 for "Structural basis for targeting human cancer antigen STEAP1 with antibodies"

### Supplementary information

#### Figures S1 – S5, Table S1

A

```
 DSTP3086S Heavy Chain       1 EVQLVESGGGLVQPGGSLRLSCAVSGYSITSDYAWNWVRQAPGKGLEWVGYISNSGSTSYNPSLKSRFTISRDTSKNTLYLQMNSLRAEDTAVYY
Fab120.545 Heavy Chain       1 DVQVQESGPGLVKPSQSLSLTCTVTGYSITSDYAWNWIRQFPGNKLEWMGYISNSGSTSYNPSLKSRISITRDTSKNQFFLQLISVTTEDTATYY
                               :**: *** ***:*. ** *:*:*:************:** **: ***:******************::*:****** ::**: *: :****.**

 DSTP3086S Heavy Chain      96 CARERNYDYDDYYYAMDYWGQGTLVTVSSASTKGPSVFPLAPSSKSTSGGTAALGCLVKDYFPEPVTVSWNSGALTSGVHTFPAVLQSSGLYSLS
Fab120.545 Heavy Chain      96 CARERNYDYDDYYYAMDYWGQGTTLTVSAASTKGPSVFPLAPSSKSTSGGTAALGCLVKDYFPEPVTVSWNSGALTSGVHTFPAVLQSSGLYSLS
                               *********************** :***:******************************************************************

 DSTP3086S Heavy Chain     191 SVVTVPSSSLGTQTYICNVNHKPSNTKVDKKVEPKSCDKTHTCPPCPAPELLGGPSVFLFPPKPKDTLMISRTPEVTCVVVDVSHEDPEVKFNWY
Fab120.545 Heavy Chain     191 SVVTVPSSSLGTQTYICNVNHKPSNTKVDKRVEPKSCDKTHTHHHHHH-----------------------------------------------
                               ******************************:***********

 DSTP3086S Heavy Chain     286 VDGVEVHNAKTKPREEQYNSTYRVVSVLTVLHQDWLNGKEYKCKVSNKALPAPIEKTISKAKGQPREPQVYTLPPSREEMTKNQVSLTCLVKGFY
Fab120.545 Heavy Chain         -----------------------------------------------------------------------------------------------

 DSTP3086S Heavy Chain     381 PSDIAVEWESNGQPENNYKTTPPVLDSDGSFFLYSKLTVDKSRWQQGNVFSCSVMHEALHNHYTQKSLSLSPGK
Fab120.545 Heavy Chain         --------------------------------------------------------------------------
```

B

```
 DSTP3086S Light Chain       1 DIQMTQSPSSLSASVGDRVTITCKSSQSLLYRSNQKNYLAWYQQKPGKAPKLLIYWASTRESGVPSRFSGSGSGTDFTLTISSLQPEDFATYYCQ
Fab120.545 Light Chain       1 DIVMSQSPSSLAVSVGEKVTMSCKSSQSLLYRSNQKNYLAWYQQKPGQSPKLLIYWASTRESGVPDRFTGSGSGTDFTLTISSVKAEDLAVYYCQ
                               ** *:******:.***::**::*************************::****************.**:**************::.**:*.****

 DSTP3086S Light Chain      96 QYYNYPRTFGQGTKVEIKRTVAAPSVFIFPPSDEQLKSGTASVVCLLNNFYPREAKVQWKVDNALQSGNSQESVTEQDSKDSTYSLSSTLTLSKA
Fab120.545 Light Chain      96 QYYNYPRTFGGGTKLEIKRTVAAPSVFIFPPSDEQLKSGTASVVCLLNNFYPREAKVQWKVDNALQSGNSQESVTEQDSKDSTYSLSSTLTLSKA
                               ********** ***:*********************************************************************************

 DSTP3086S Light Chain     191 DYEKHKVYACEVTHQGLSSPVTKSFNRGEC
Fab120.545 Light Chain     191 DYEKHKVYACEVTHQGLSSPVTKSFNRGEC
                               ******************************
```

**Figure S1:** Amino-acid sequence alignment between the heavy (A) and light (B) chains of clinically employed antibody DSTP3086S and the antibody-fragment Fab120.545 used in our work. The CDR regions are boxed and colored yellow. Amino acids that contribute to the binding to STEAP1 are marked bold. The hexa-His tag at the C-terminus of Fab120.545, which is used for purification, is colored blue.

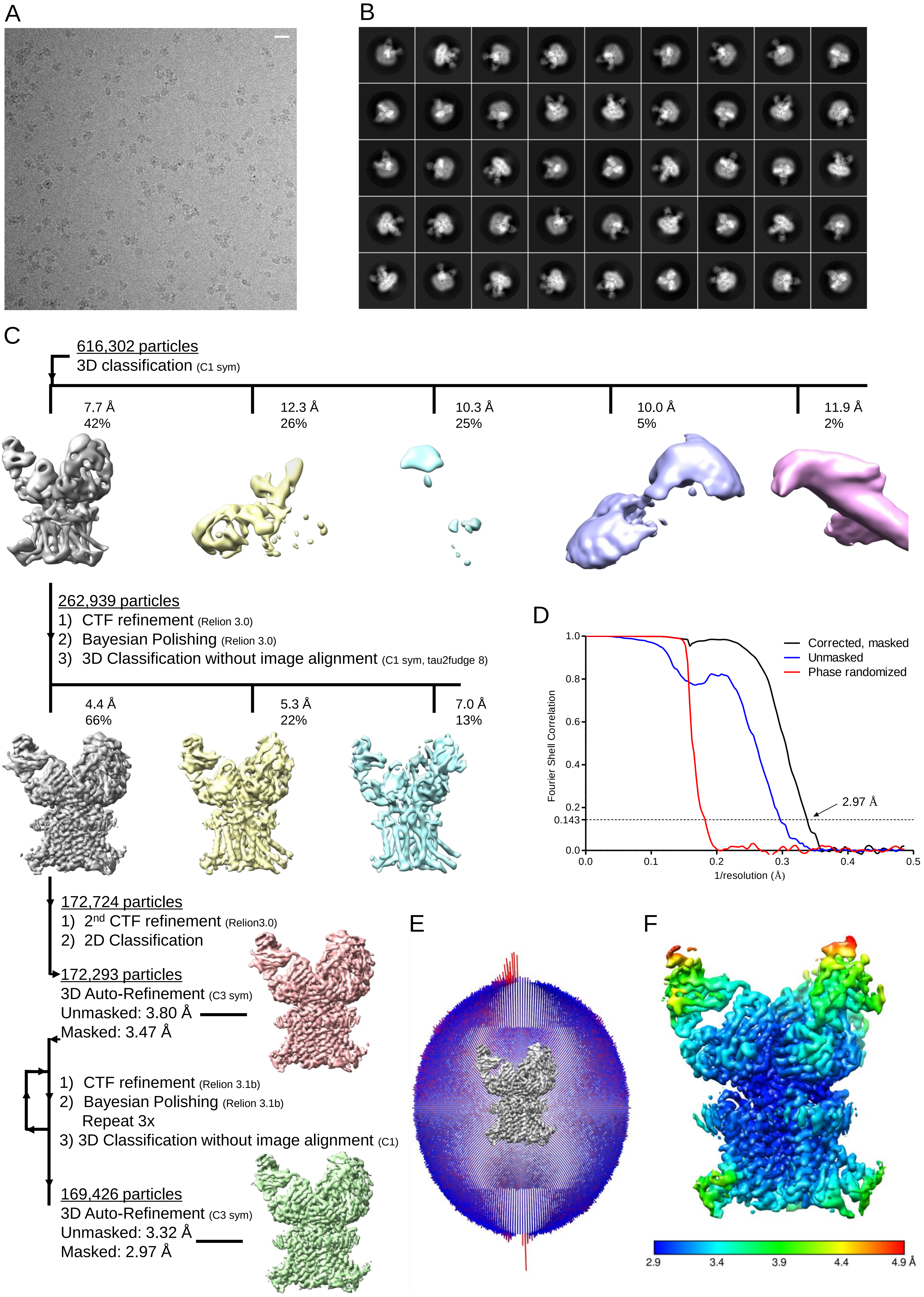


**Figure S2:** Cryo-EM image processing. (A) Micrograph depicting STEAP1-Fab120.545 particles distributed in vitreous ice. The scale bar length is 200 Å. (B) Exemplary 2D-class averages generated in Relion. (C) Image processing strategy in the Relion pipeline. All shown density maps are unsharpened and are depicted in the same orientation. (D) Fourier shell correlation plot for gold-standard refined masked (black), unmasked (blue) and high-resolution phase randomized (red) half maps. The FSC = 0.143 threshold is shown as a dashed line. (E) Angular distribution of the particles that were used to reconstruct the final STEAP1-Fab120.545 density map with C3 symmetry applied. (F) Local-resolution estimation of the reconstruction, computed through Relion.

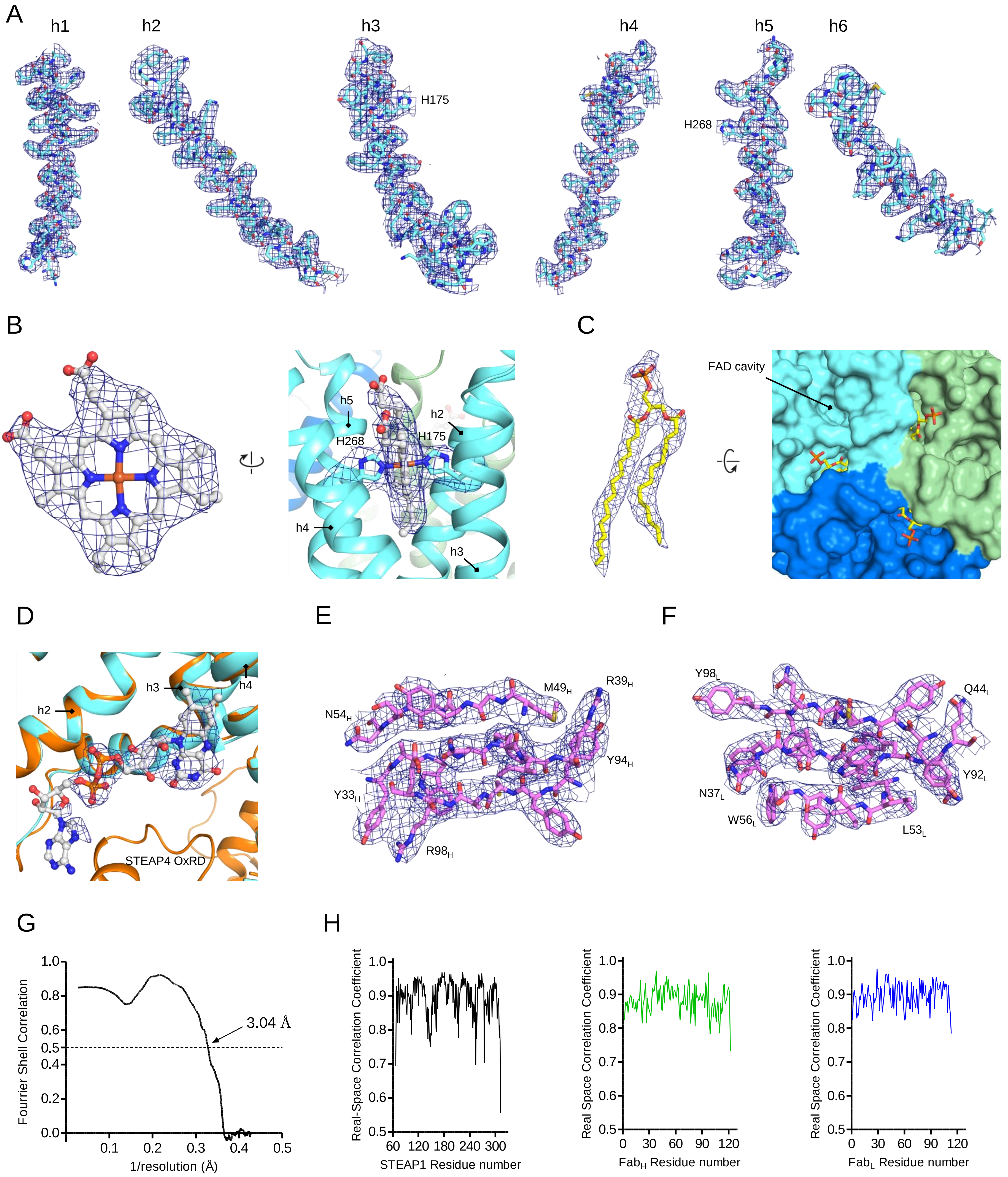


**Figure S3:** Modelling of the STEAP1-Fab120.545 structure and model validation. (A) Cryo-EM density of the six membrane helices of STEAP1 with modelled amino acid residues shown as sticks. Annotated residues H175 and H268 coordinate the central-heme iron. (B) Isolated density with fitted model of the heme cofactor (left panel), and the protein environment of the heme-binding site at the extracellular side of the membrane (right panel). (C) Isolated density with fitted model of a lipid molecule (left panel). The lipid was modelled as 14:0 Phosphatidic acid (DMPA), but the observed density could represent a mixture of several partially bound lipids. The right panel shows the packing of the lipid in cavities between STEAP1 subunits (depicted as surface) orthogonal to the membrane from the cytoplasmic side. (D) Overlay of STEAP1 and STEAP4 structures at the intracellular side of the TMD. The FAD cofactor modelled in the STEAP4 structure is shown in ball-and-stick representation, while non-protein residue density of the STEAP1 map is depicted as mesh. The position of the FAD model was not changed after rigid-body fitting STEAP4 on the STEAP1 structure. Helix h5 resides in front of FAD and is omitted from the figure for clarity. (E) Selected Beta-sheet density with fitted amino acids of the Fab120.545 heavy chain. (F) Selected Beta-sheet density with fitted amino acids of the Fab120.545 light chain. (G) Fourier shell correlation plot of the final reconstructed map versus the build STEAP1-Fab model as determined by Phenix. The FSC = 0.5 threshold is shown as a dashed line. (H) Real-space correlation coefficient plotted for every amino-acid residue as calculated by Phenix for STEAP1 (left panel) and Fab heavy (middle panel) and light (right panel) chains.

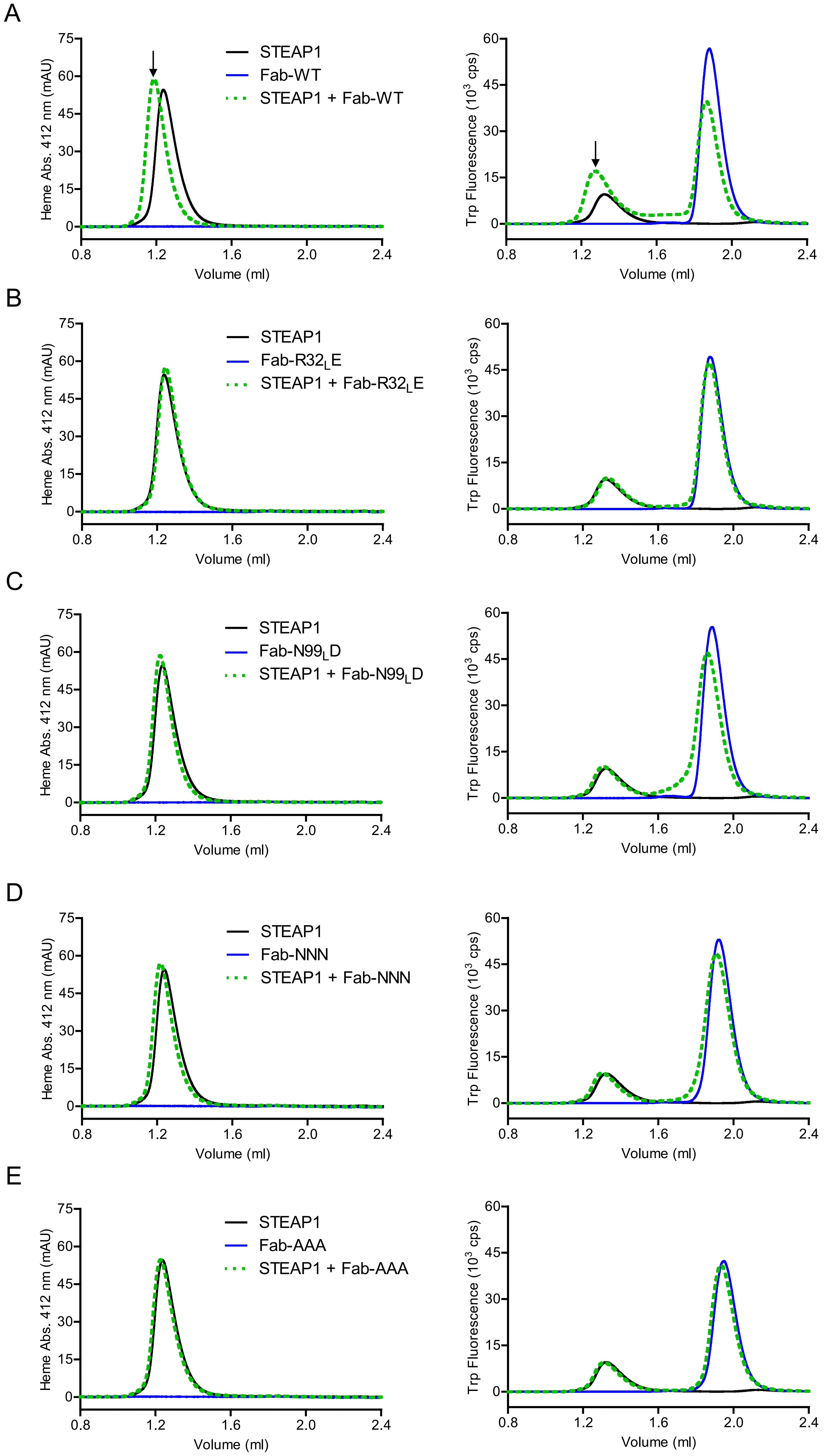


**Figure S4:** Size-exclusion chromatography-based assays to test the interaction of STEAP1 with (A) wildtype, (B) R32$_L$E, (C) N99$_L$D, (D) NNN (D103$_H$N, D105$_H$N, D106$_H$N), (E) AAA (D103$_H$A, D105$_H$A, D106$_H$A) variants of Fab120.545. The binding event in panel A (annotated with an arrow) is established through a peak shift in the heme-absorbance profile, and a peak shift and a complex peak-height increase in the Trp-fluorescence profile. None of the tested mutants (panels B – E) show clear binding to STEAP1.

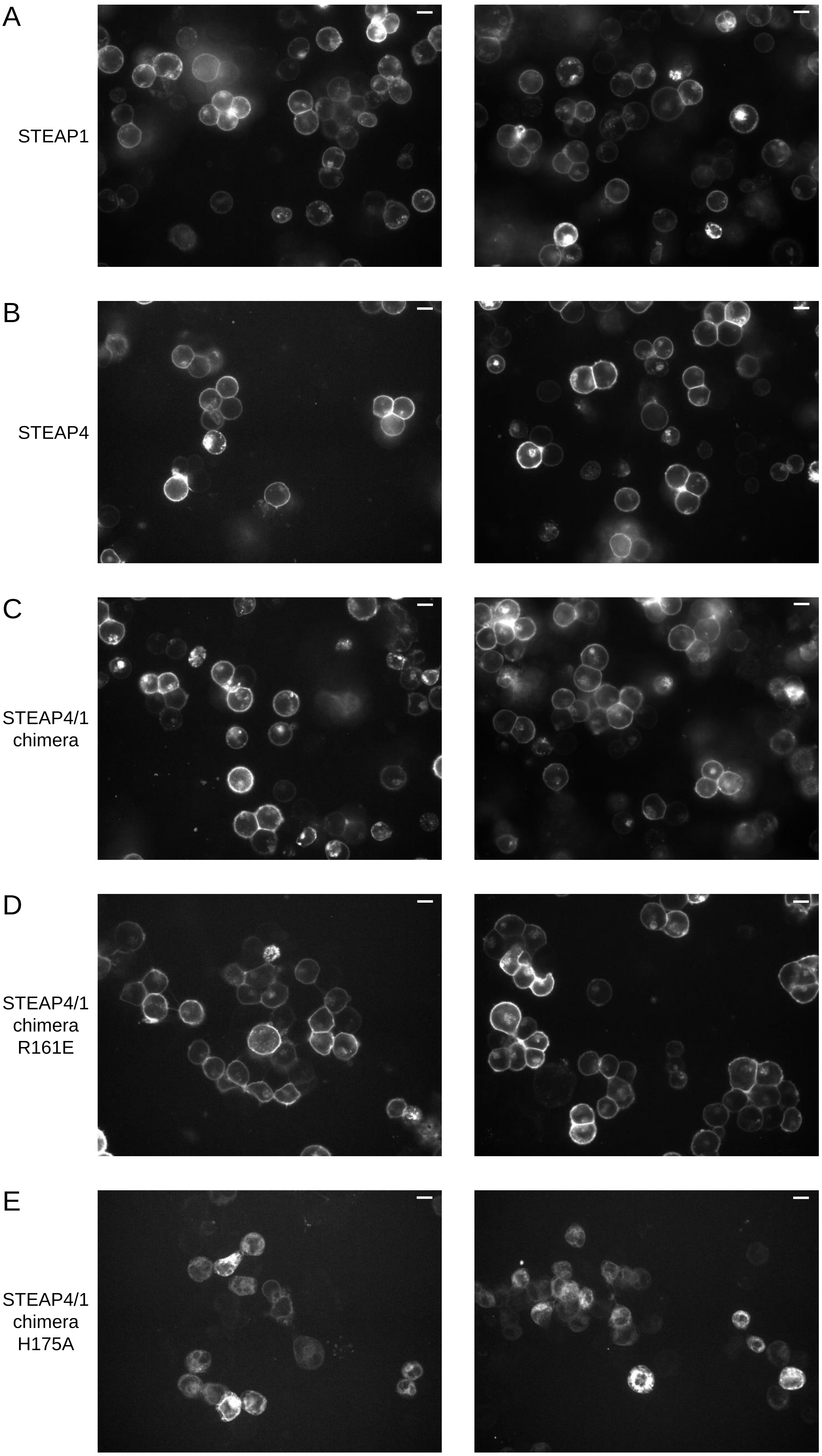


**Figure S5:** Confocal microscopy of HEK293 GNT1$^-$ suspension cells expressing GFP-tagged STEAP variants. Two representative images are shown for cells expressing (A) STEAP1, (B) STEAP4, (C) STEAP4/1$_{chimera}$, (D) STEAP4/1$_{chimera}$-R161E, (E) STEAP4/1$_{chimera}$-H175A. The only protein that did not (partially) localize to the plasma membrane was STEAP4/1$_{chimera}$-H175A (panel E). The scale bar length is 10 µm.

#### Table S1. Cloning and mutagenesis primers for STEAP constructs

| **Cloning primers** | | | |
|---|---|---|---|
| Construct | Direction | Residue # | Sequence |
| STEAP4/$1_{chimera}$ (STEAP4 Gibson) | Forward | 1 | GTCCAGAGCTCGGATCCGAGAAAACCTGCATCGACGCCCTGC |
| STEAP4/$1_{chimera}$ (STEAP4 Gibson) | Reverse | 198 | GCAGATGCCACTGGGGGAACAGCTGCAGGGGG |
| STEAP4/$1_{chimera}$ (STEAP1 Gibson) | Forward | 67 | CCCCCTGCAGCTGTTCCCCCAGTGGCATCTGC |
| STEAP4/$1_{chimera}$ (STEAP1 Gibson) | Reverse | 339 | CGATGCAGGTTTTCTCGGATCCGAGCTCTGGACAAGACACGTGGC |

| **Mutagenesis primers** | | |
|---|---|---|
| Construct | Direction | Sequence |
| STEAP4/$1_{chimera}$-R161E | Forward | GATGCTGACCGAGAAGCAGTTCGG |
| STEAP4/$1_{chimera}$-R161E | Reverse | CACTTGTCCAGCCAGTGA |
| STEAP4/$1_{chimera}$-H175A | Forward | CGCCGTGCTGGCCGCCATCTACA |
| STEAP4/$1_{chimera}$-H175A | Reverse | AAGAAGAAGCTCAGCAGG |
